# Integrative analysis of microbial 16S gene and shotgun metagenomic sequencing data improves statistical efficiency

**DOI:** 10.1101/2023.06.27.546795

**Authors:** Ye Yue, Timothy D. Read, Veronika Fedirko, Glen A. Satten, Yi-Juan Hu

## Abstract

The most widely used technologies for profiling microbial communities are 16S marker-gene sequencing and shotgun metagenomic sequencing. Interestingly, many microbiome studies have performed both sequencing experiments on the same cohort of samples. The two sequencing datasets often reveal consistent patterns of microbial signatures, highlighting the potential for an integrative analysis to improve power of testing these signatures. However, differential experimental biases, partially overlapping samples, and differential library sizes pose tremendous challenges when combining the two datasets. Currently, researchers either discard one dataset entirely or use different datasets for different objectives. In this article, we introduce the first method of this kind, named Com-2seq, that combines the two sequencing datasets for the objective of testing differential abundance at the genus and community levels while overcoming these difficulties. We demonstrate that Com-2seq substantially improves statistical efficiency over analysis of either dataset alone and works better than two *ad hoc* approaches.

---

Thanks to technological advances in high-throughput sequencing, microbiome research has proliferated in the past decade and revealed relationships between human microbiome and many diseases and conditions such as inflammatory bowl diseases [1], obesity and type II diabetes [2], and even cancers [3]. The most widely used technologies for profiling microbial communities are 16S marker-gene sequencing (16S) and shotgun metagenomic sequencing (SMS) [4]. The 16S method employs primers that target a highly variable region of the 16S ribosomal RNA gene, which is then PCR amplified and sequenced. This approach is well-tested, fast and cost effective, and provides a low-resolution view of a microbial community, typically at the genus level. On the other hand, SMS extracts all microbial genomes within a sample, which are then fragmented and sequenced. This technique offers detailed genomic information, including higher taxonomic resolution and additional functional capability, but it is 10 to 30 times more expensive to perform sequencing experiment and much more challenging to conduct bioinformatics analysis.

Both the 16S and SMS methods are subject to *experimental bias* at every step of the experiment (i.e., DNA extraction, PCR amplification, amplicon or metagenomic sequencing, and bioinformatics processing), as each step favors certain taxa over others [5]. The bias systematically distorts the measured taxon abundances from their actual values. Moreover, the bias differs significantly between 16S and SMS, as they comprise different steps (e.g., PCR amplification vs. no PCR) and even for the same step they may adopt different protocols [6]. As a result, each sequencing method leads to some taxa being underrepresented or even entirely missed [7]. As these taxa may be complementary, combining data from the two sequencing experiments could enhance the profiling of complex microbial communities.

Interestingly, many microbiome studies have performed both 16S and SMS on the same cohort of samples. In Qiita, which is currently the largest open-source microbiome study management platform (https://qiita.ucsd.edu) [8], 26 out of all 698 studies (as of June 2023) have both datasets. In fact, more studies may have both datasets, as some studies may only deposit the dataset used in the final publication. There are at least two scenarios in which this occurs. In one scenario, a study initially performs 16S and then follows up with SMS to investigate higher-resolution taxa or biological functions. In another scenario, a study initially performs SMS but adds 16S due to its low cost. The 16S data are routinely processed into a taxa count table by the popular analysis platform QIIME2 [9]. Although there is less consensus, the SMS data can also be classified into taxonomies using tools such as MetaPhlAn [10–12], Kraken [13, 14], or more recently Woltka [15]. For each of the 26 studies found in Qiita, we summarized the two taxa count tables, as well as their overlapping samples and taxa at the genus level, in Table S1. For most studies, both the overlap of samples and the overlap of genera are incomplete yet substantial, which data structure is schematically depicted in Figure 1 (left). The library sizes (i.e., depths) in the two tables may differ by orders of magnitude, ranging from 1.4 to 1500 fold. Despite their differences, the 16S and SMS taxonomic profiles for the same cohort of samples have frequently been found to yield consistent patterns of microbial signatures [7, 16–22]. This further underscores the need for an integrative analysis of the two taxonomic profiles to improve power of testing these signatures.

**Figure 1:**
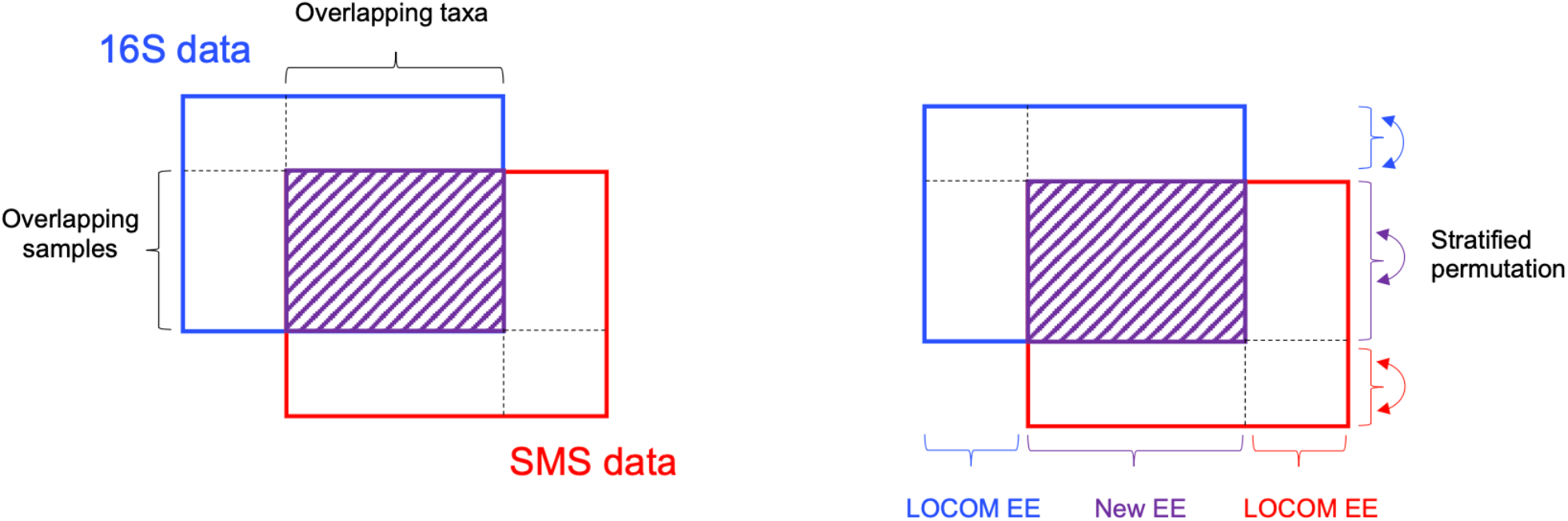
Illustration of the data structure (left) and strategies for analyzing the data (right). The shaded area indicates the data for the overlapping samples and taxa between the 16S and SMS taxa count data. New EE is Equation (1) or bias-corrected Equation (S3) in this paper. LOCOM EE is the estimating equation in the LOCOM paper [23].

Currently, researchers either discard one dataset entirely or use different datasets for different objectives. To the best of our knowledge, no statistical method is available for performing integrative analysis of the two sequencing datasets. Therefore, we propose a method, named Com-2seq, to fill this gap. Our method is based on same model that underlies the LOCOM package we developed recently [23], which employs logistic regression for testing taxon differential abundance while remaining robust to experimental bias. LOCOM is a compositionally aware method that does not require pseudocounts. Our new method combines data from 16S and SMS for testing differential abundance at the genus and community levels, while overcoming the challenges including differential experimental biases, partially overlapping samples, and differential library sizes. We compare our new method to two *ad hoc* procedures: one first pools the taxon counts of the two datasets and then applies LOCOM to the pooled data, referred to as Com-count; the other applies LOCOM to the two datasets separately and then combines the two *p*-values at each taxon using the Cauchy *p*-value combination method, referred to as Com-p; see Methods for more details.

## Results

### Differences in experimental bias

We compared the observed relative abundances between 16S and SMS for each of the overlapping samples at each of the overlapping genera, using the ORIGINS and Dietary study data from Qiita; descriptions about the two studies are provided below. If the biases in 16S and SMS data were the same, we would expect data from each genus to fit the 45° line. The departure from this behavior seen in Figure 2 for the ORIGINS study demonstrates that there are systematic differences in observed relative abundances at many genera, even for the second most abundant genus *Haemophilus*. Further, some genera appear to be mostly absent in one dataset but reasonably abundant in the other, such as *Gemella* from 16S and *Geobacillus* from SMS. These findings confirm that 16S and SMS data have different experimental biases. Similar patterns, although with higher overdispersion (i.e., deviation of data from the fitted line), are observed in Figure 3 for the Dietary study. Despite the differences, there is clear agreement between 16S and SMS data, particularly at the most abundant genera from the ORIGINS study, motivating the use of an integrative approach that properly accounts for the differences in experimental bias.

**Figure 2:**
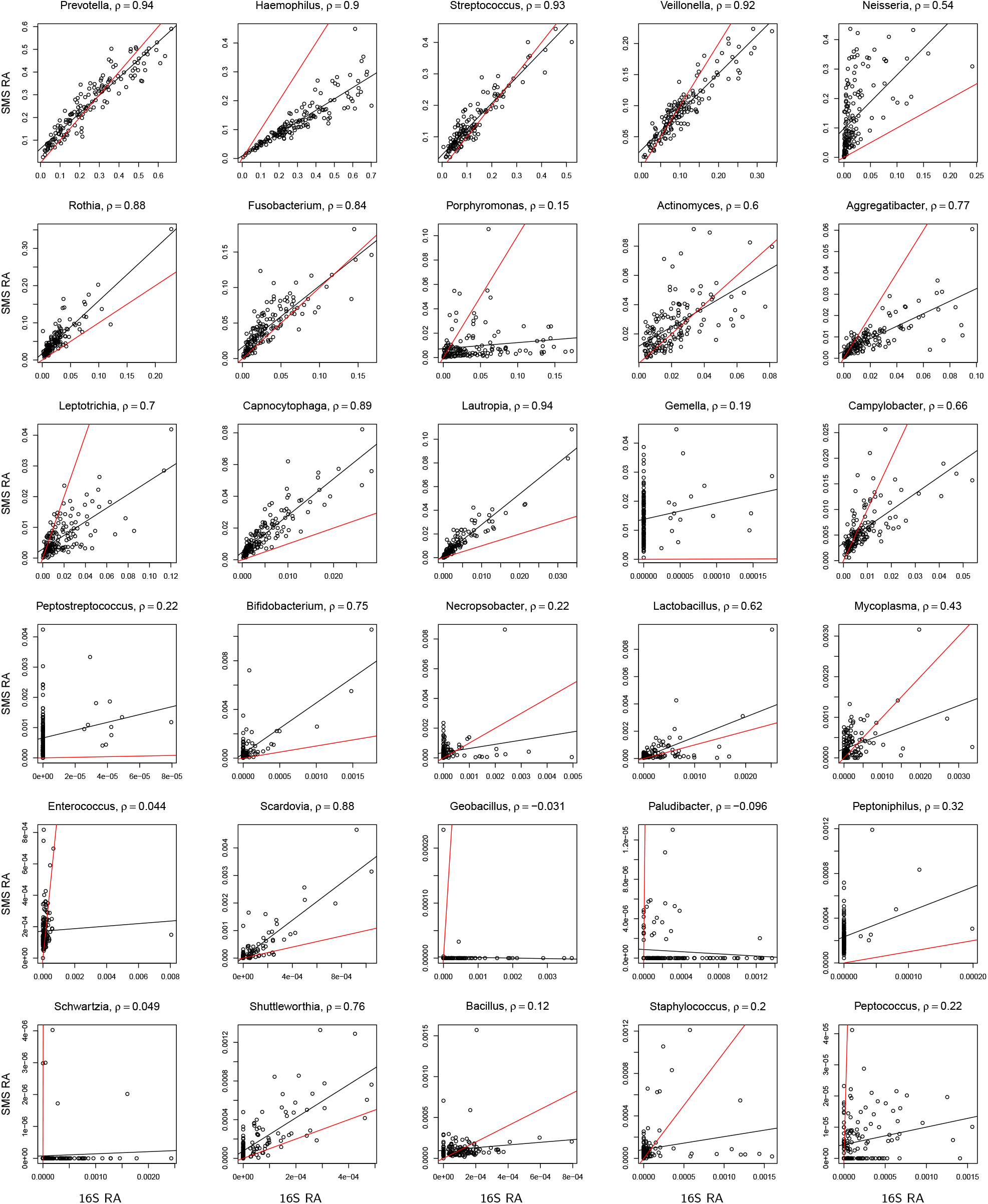
Scatter plot of observed relative abundances (RA) between 16S and SMS for the top 1–15 and 36–50 most abundant genera (ordered by decreasing abundance) in the ORIGINS study. The *ρ* value is the Pearson correlation coefficient. The red line is the 45^*°*^ reference line. The black line depicts a fitted linear regression. The observed relative abundances were calculated based on the 125 overlapping genera.

**Figure 3:**
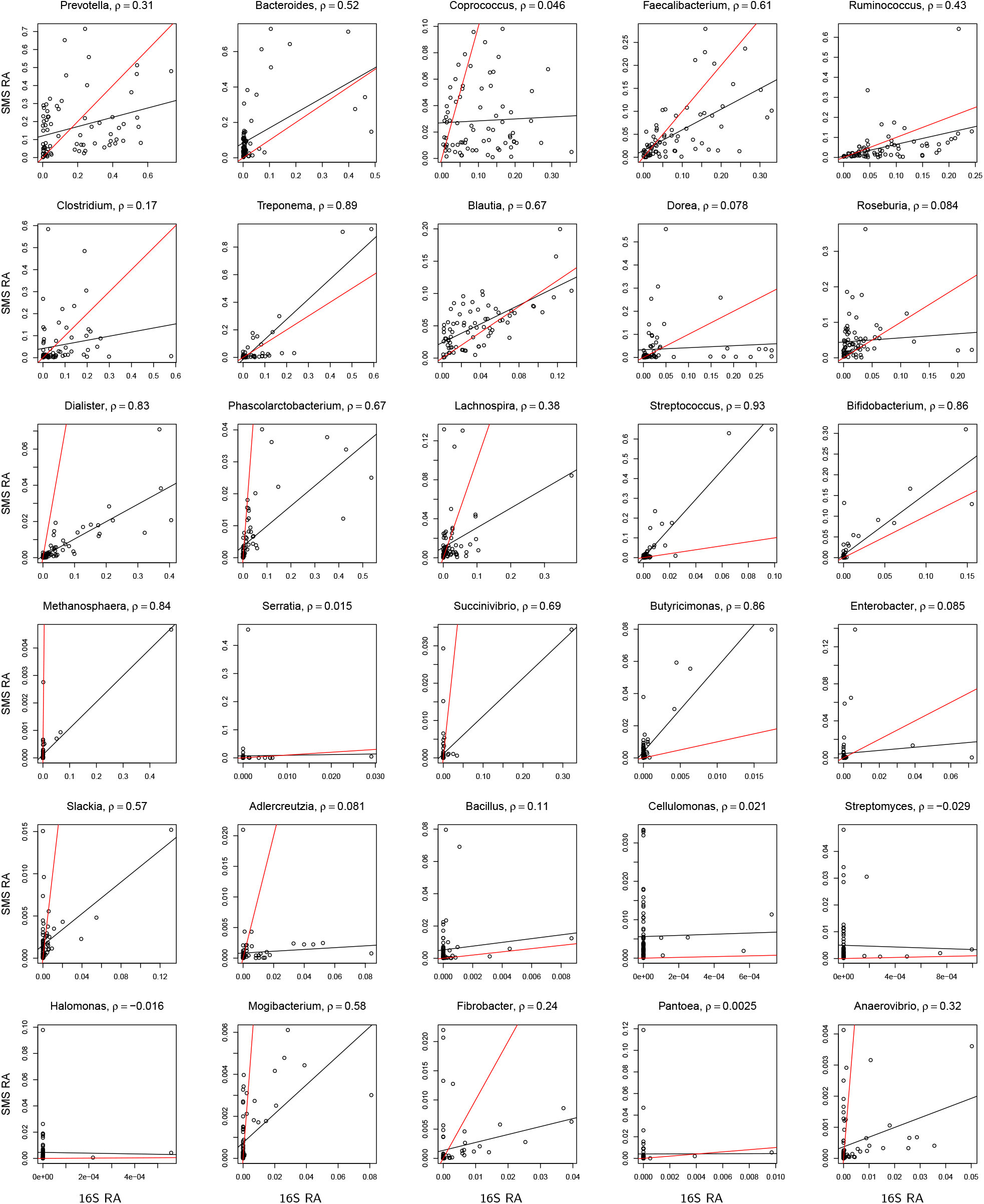
Scatter plot of observed relative abundances between 16S and SMS for the top 1–15 and 36–50 most abundant genera in the Dietary study. See more information in the caption of Figure 2.

### Analysis of ORIGINS data

We analyzed data generated from the Oral Infections, Glucose Intolerance, and Insulin Resistance Study (ORIGINS) [24] to investigate the association between periodontal bacteria and prediabetes status (yes or no) among diabetes-free adults, without adjusting for any other risk factors. One subgingival plaque sample was collected from each participant and sequenced by either 16S, SMS, or both. We downloaded the 16S and SMS taxa count data with study ID 11808 from Qiita. The 16S data include 271 samples linked with meta data and having adequate (*>* 5,000) library sizes, and 234 genera after quality control (QC, which excluded genera found in fewer than 5 samples). The SMS data include 183 samples linked with meta data and having adequate library size, and 756 genera after QC. In total, there are 302 distinct samples (56 cases and 246 controls) and 864 distinct genera, among which 152 samples (47 cases and 105 controls) have both 16S and SMS data for 125 genera. The mean library sizes of the 16S and SMS data are 26,950 and 176,321, respectively, resulting in a depth ratio of 1:6.5.

We began by analyzing data for the overlapping samples and genera, with a main purpose of method comparison. We applied Com-2seq, Com-count, and Com-p for integrative analysis of the 16S and SMS data, as well as LOCOM to analyze each dataset separately, and summarized their global test *p*-values and detected differentially abundant genera (at the nominal FDR level of 10%) in Table 1. Com-2seq and Com-count yielded significant global *p*-values (at the nominal level of 0.05) and detected 7 and 3 genera, respectively, whereas the analysis of either dataset alone and the integrative analysis by combining *p*-values (Com-p) produced non-significant global *p*-values and detected zero or at most one genus. For more information on the detected genera, see Table S2 and Figure S1.

**Table 1:**
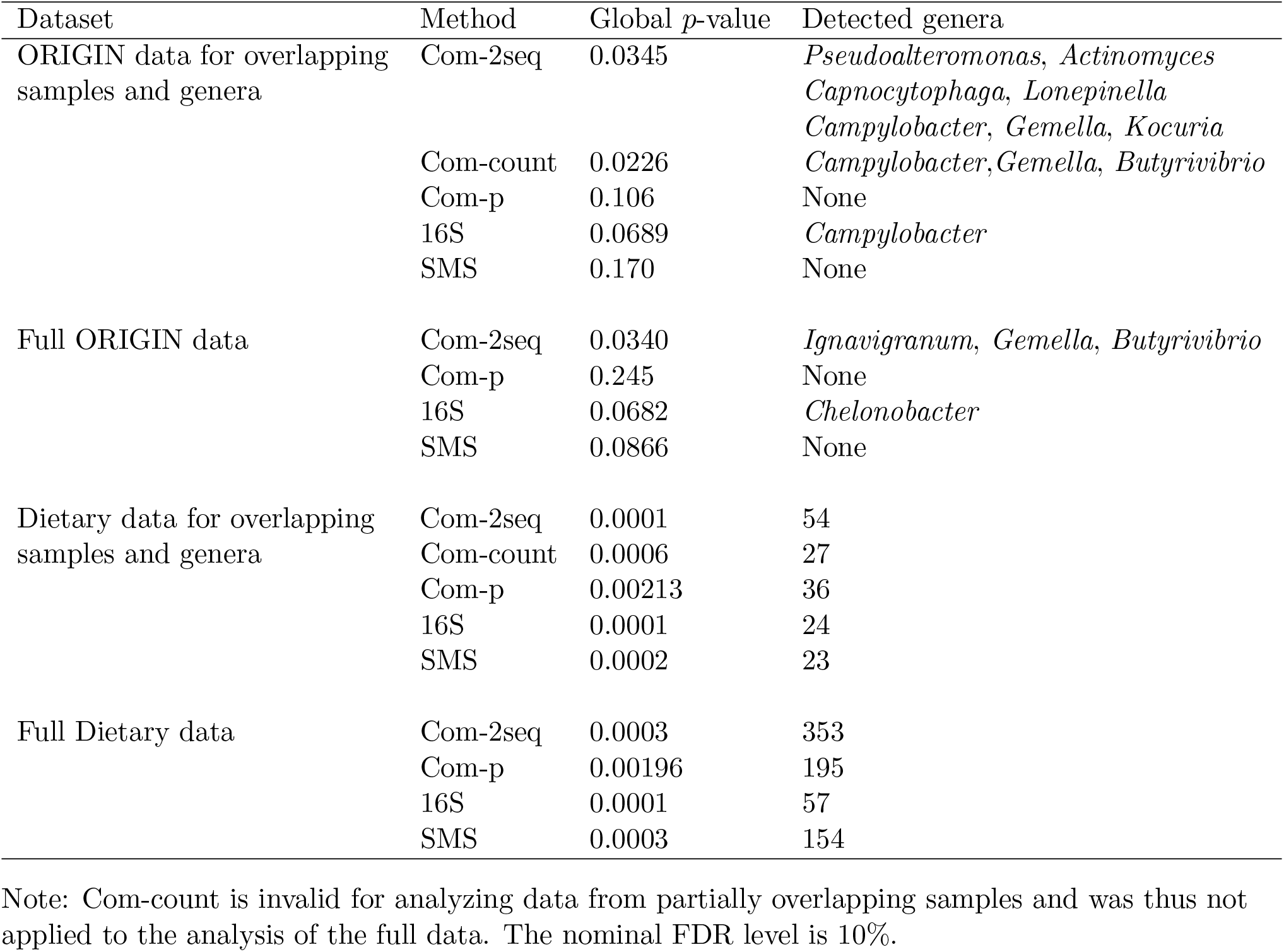
Global test *p*-value and detected differentially abundant genera in the analysis of the real datasets

| Dataset | Method | Global $p$-value | Detected genera |
| --- | --- | --- | --- |
| ORIGIN data for overlapping samples and genera | Com-2seq | 0.0345 | *Pseudoalteromonas*, *Actinomyces*<br>*Capnocytophaga*, *Lonepinella*<br>*Campylobacter*, *Gemella*, *Kocuria* |
|  | Com-count | 0.0226 | *Campylobacter*, *Gemella*, *Butyrivibrio* |
|  | Com-p | 0.106 | None |
|  | 16S | 0.0689 | *Campylobacter* |
|  | SMS | 0.170 | None |
| Full ORIGIN data | Com-2seq | 0.0340 | *Ignavigranum*, *Gemella*, *Butyrivibrio* |
|  | Com-p | 0.245 | None |
|  | 16S | 0.0682 | *Chelonobacter* |
|  | SMS | 0.0866 | None |
| Dietary data for overlapping samples and genera | Com-2seq | 0.0001 | 54 |
|  | Com-count | 0.0006 | 27 |
|  | Com-p | 0.00213 | 36 |
|  | 16S | 0.0001 | 24 |
|  | SMS | 0.0002 | 23 |
| Full Dietary data | Com-2seq | 0.0003 | 353 |
|  | Com-p | 0.00196 | 195 |
|  | 16S | 0.0001 | 57 |
|  | SMS | 0.0003 | 154 |

Note: Com-count is invalid for analyzing data from partially overlapping samples and was thus not applied to the analysis of the full data. The nominal FDR level is 10%.

We proceeded to analyze the full data for scientific discovery and also summarized the main results in Table 1. In this case, Com-count is invalid (as demonstrated in simulation results below) and was thus not applied. Once again, Com-2seq yielded a significant global *p*-value, while the *p*-values of the remaining methods failed to reach the significance level. Com-2seq detected three genera, *Butyrivibrio, Gemella*, and *Ignavigranum*, whereas the analysis of the 16S data alone detected a different genus *Chelonobacter* and the analysis of the SMS data alone and the integrative analysis by Com-p failed to detect any genus. More information on the detected genera can be found in Figure 4(a) and Table S3. Specifically, *Butyrivibrio* was captured by both sequencing techniques and assigned non-significant yet small *p*-values in both analyses of individual datasets. As its abundance appeared consistently lower in prediabetic participants across both datasets, this trend was deemed significant overall by Com-2seq. Indeed, *Butyrivibrio* is known to produce short-chain fatty acids such as butyrate, which has been found to improve insulin sensitivity in mice [25]. *Gemella* was only identified in a few samples by 16S sequencing and thus excluded from the analysis of 16S data alone by the LOCOM filter. On the other hand, it was efficiently captured in all samples by SMS (*∼*1.5% relative abundance) and revealed to be notably more abundant in prediabetic participants, resulting in its detection by Com-2seq (which analysis salvaged the 16S data on this genus that were discarded in the analysis of 16S data alone). This finding is also plausible, as *Gemella* has been linked to various infections, including those affecting heart valves [26], brain membranes [27], and bloodstreams [28]. *Ignavigranum* was completely missed by 16S sequencing and its detection by Com-2seq was solely driven by its differential abundance revealed by SMS. *Chelonobacter* exhibited significant differential abundance using the 16S data, but this signal was not replicated using the SMS data, leading to a non-significant overall result by Com-2seq.

**Figure 4:**
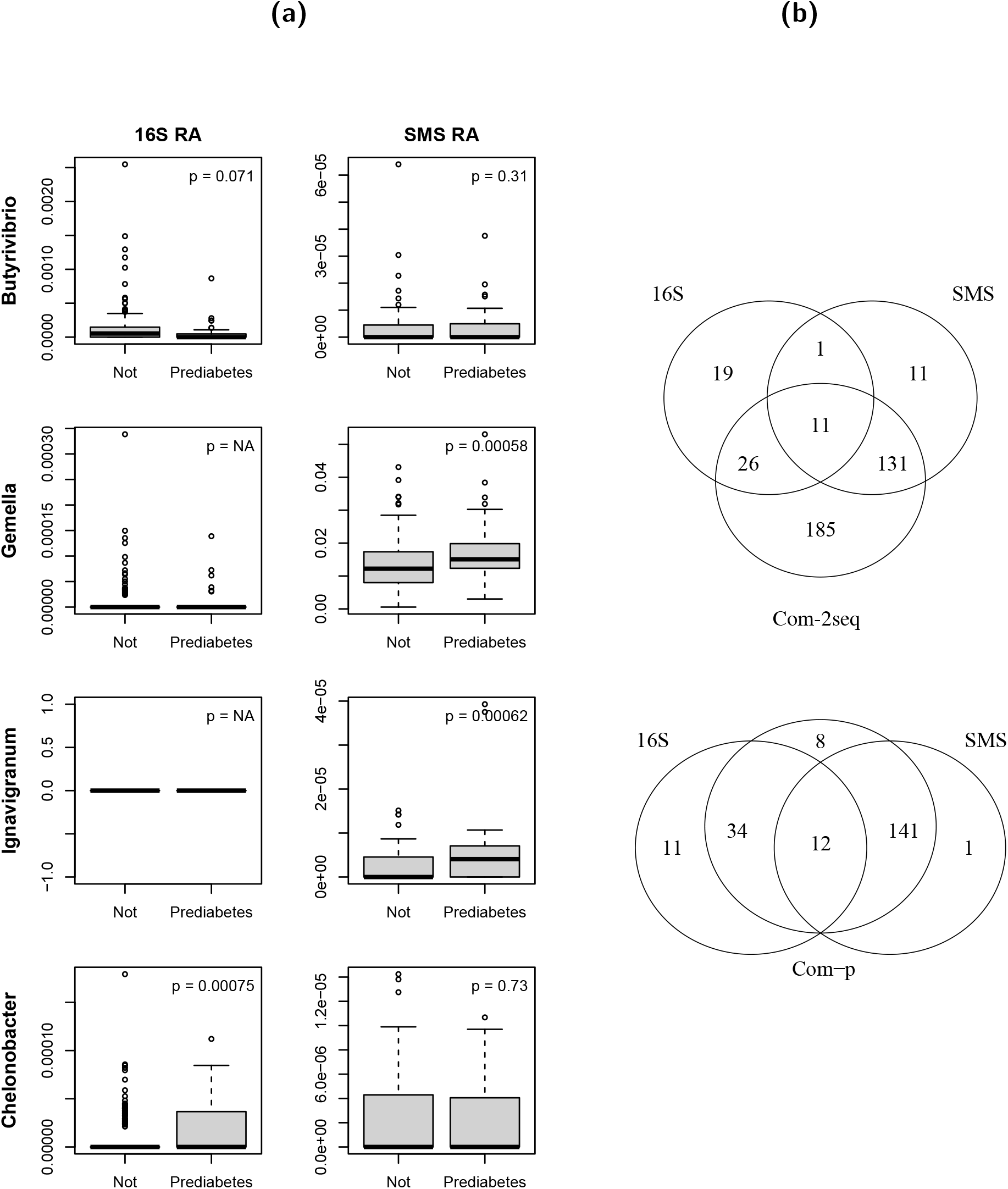
More information on the detected genera in the analysis of the real datasets. (a) Distribution of observed relative abundances (RA) by prediabetes status for the detected genera in the analysis of the full ORIGINS data. The observed relative abundances were calculated based on all genera for all samples in the 16S or SMS data. The *p*-values are from the analysis of individual 16S and SMS datasets, as also listed in Table S3. (b) Venn diagram for the detected genera in the analysis of the full Dietary data.

### Analysis of Dietary data

We also analyzed data from Amato et al. [29] to test the influence of host dietary niche (folivore vs. non-folivore) on the gut microbiome of wild non-human primates, while controlling for host phylogeny (categorized as apes, lemurs, new world monkeys, and old world monkeys). One fecal sample was collected for each animal and sequenced by either 16S, SMS, or both. We downloaded the 16S and SMS taxa count data with study ID 11212 from Qiita, the statistics of which are listed in Table S1. In total, there are 172 distinct samples (94 folivore and 78 non-folivore) and 2062 distinct genera, among which 76 samples (40 folivore and 36 non-folivore) have both 16S and SMS data for 236 genera. The ratio of 16S to SMS mean library sizes is 1:9.8.

Table 1 (lower panel) shows that all analysis methods yielded highly significant global *p*-values and detected a large number of differentially abundant genera at the nominal FDR level of 10%, in both the analysis of data for the overlapping samples and genera and the analysis of the full data. In both of these analyses, Com-2seq detected the most genera, far exceeding the number of detections by analyzing the 16S or SMS data alone or by employing Com-p. Figure 4(b) displays the Venn diagram of the detected genera by different methods when analyzing the full data. Com-2seq detected 185 novel genera that were missed by analyzing the 16S and SMS data separately, whereas Com-p detected only 8 such genera.

### Simulation results

To develop confidence in our new method, we used simulated data to confirm the results we observed in the analysis of the ORIGINS and Dietary data; the details of the simulation studies are presented in Methods. In brief, we considered three causal mechanisms M1, M2, and M3, which assume the causal (i.e., trait-associated) taxa to be moderately abundant, very abundant, and rare, respectively. We varied the overdispersion parameter *τ*, which controls the deviation of 16S and SMS relative abundances from the fitted line (e.g., Figures 2 and 3), between 0.01 and 0.001, and we also varied the ratio of 16S to SMS mean library sizes between 1:1 and 1:10.

The results for the complete-overlap case, in which all samples have both 16S and SMS data, based on each of the four combinations of the overdispersion and depth ratio values are shown in Figures 5 and S2–S4, respectively. All global tests controlled the type I error at the nominal level and all taxon-level tests controlled the FDR at the nominal level across all scenarios. All integrative analyses significantly improved statistical efficiency over the analyses of individual datasets, at both the taxon and global levels. The increase in sensitivity of detecting causal taxa is substantial, as one sequencing experiment failed to capture certain causal taxa and the combination of two experiments provided better coverage. The boost in global power is less pronounced, because the Harmonic Mean and Cauchy statistics underlying the global tests were dominated by a few smallest individual *p*-values and less influenced by the total number of causal taxa. Among the three integrative approaches, Com-2seq always yielded the highest or nearly highest power and sensitivity.

**Figure 5:**
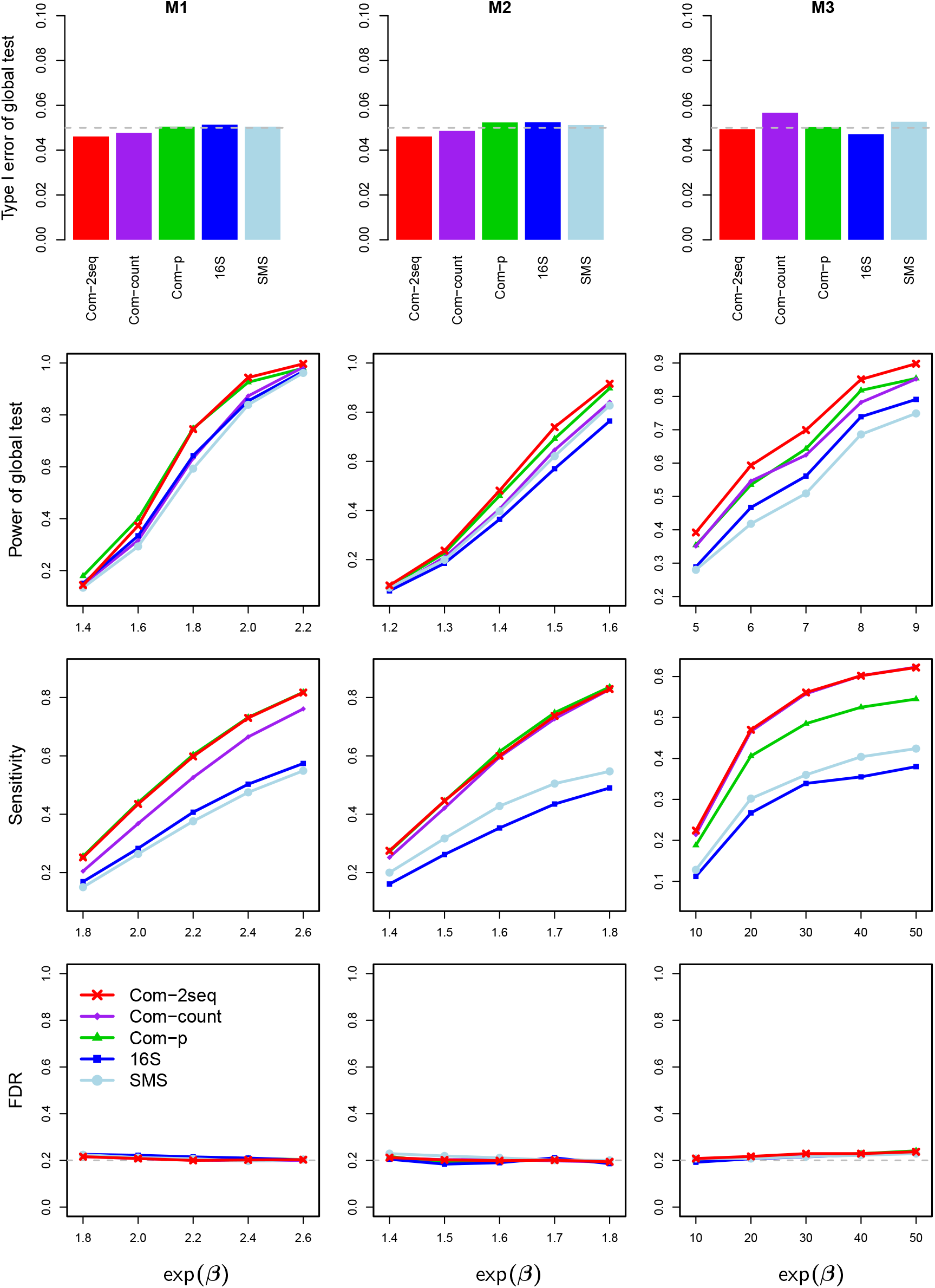
Type I error rate (first row, when *β* = 0) and power (second row) for testing the global association at the nominal level of 0.05 (gray dashed line), and sensitivity (third row) and empirical FDR (fourth row) for testing individual taxa at the nominal FDR level of 0.2 (gray dashed line), based on data simulated with completely overlapping samples, overdispersion of *τ* = 0.01, and depth ratio of 1:10.

Similar patterns of results are observed in the partial-overlap case (Figure 6), in which part of samples have both 16S and SMS data and the remaining samples have data from either 16S or SMS. One difference from the complete-overlap case is that, Com-count failed to control the type I error, as its zero-filling strategy tended to create spurious associations. Another difference pertains to the permutation scheme in Com-2seq, which was based on three strata of samples in the partial-overlap case, as illustrated in Figure 1 (right). It is the only one that led to correct type I error, compared to two alternative schemes: one pools samples having data from either of the two experiments into one stratum (a total of two strata), and the other pools all samples together (one stratum). Finally, the same patterns of results persisted when a confounder was simulated (Figure S5). We confirmed that the confounding effect was substantial, leading to highly inflated type I error if it were not controlled for. All methods yielded proper type I error after adjusting for the confounder.

**Figure 6:**
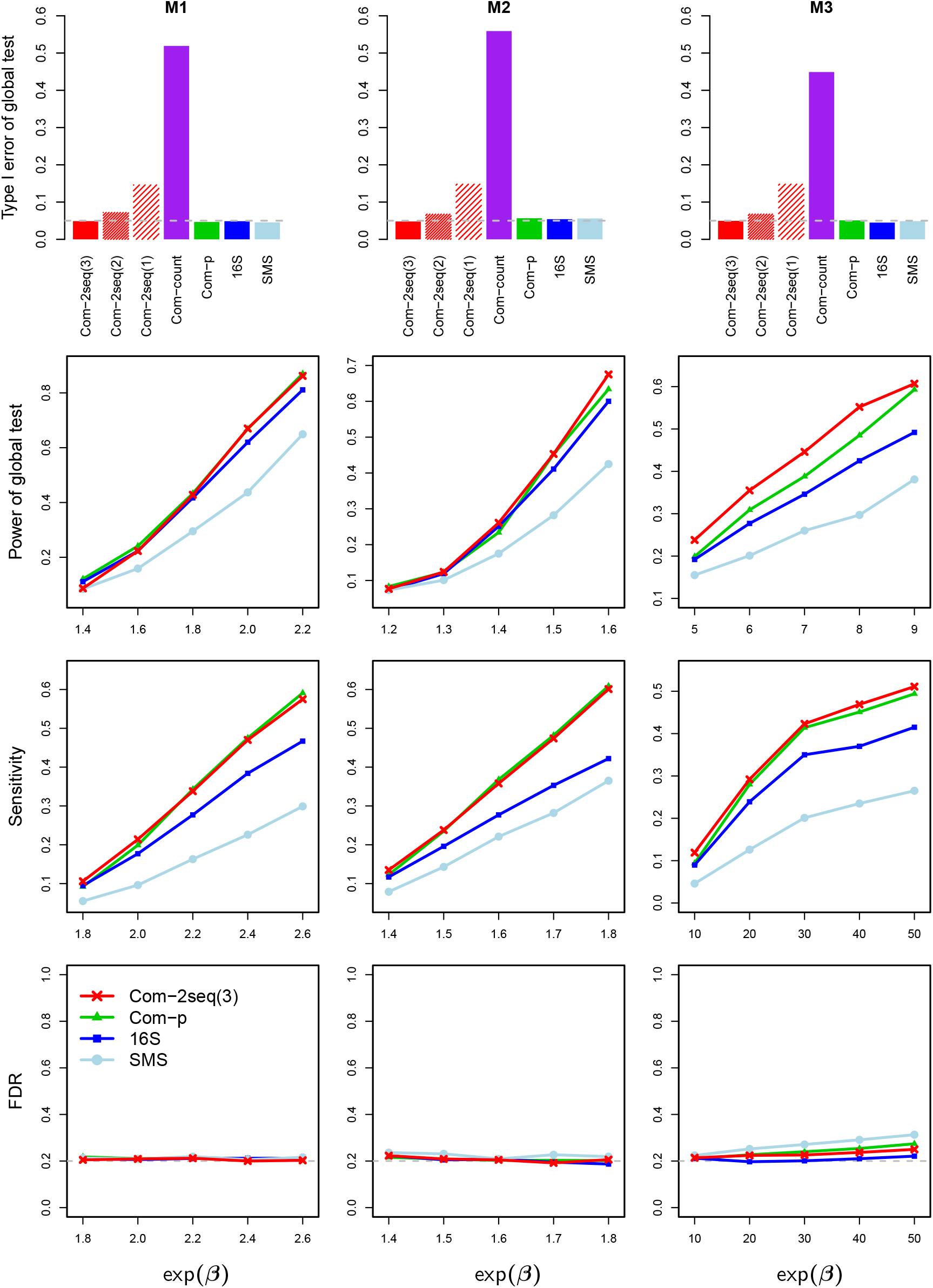
Results for data simulated with partially overlapping samples. Com-2seq annotated with “(3)”, “(2)” and “(1)” used permutation schemes that were based on three strata (the proposed one), two strata, and one stratum, respectively. See more information in the caption of Figure 5.

## Discussion

We have presented a new method, Com-2seq, that is the first method for integrative analysis of 16S and SMS data. Com-2seq is specifically designed to combine these datasets in a way that properly accounts for their differences in experimental bias, sequenced samples, and library sizes. We have demonstrated that Com-2seq substantially improves statistical efficiency over analysis of either of the two datasets and works better than the taxa-count-pooling and *p*-value-combination approaches. The inference of Com-2seq is based on a permutation procedure that preserves any sample structure and thus valid for partially overlapping samples as well as small sample sizes. Com-2seq allows testing of a trait that is binary, continuous, or multivariate (e.g., a categorical trait with more than two levels or a set of multiple measurements), and supports adjustment of confounding covariates.

We have implemented Com-2seq as a new function in our R package LOCOM, which is available on GitHub at https://github.com/yijuanhu/LOCOM. Com-2seq performs best when given count data. However, some bioinformatics programs for processing shotgun metagenomic data, such as Kraken, output relative abundance data only. In this case, Com-2seq can still be used, although the performance may be slightly worse than if count data were available. Com-2seq automatically adjusts to handle only relative abundance data.

## Methods

### Com-2seq

Let *Y*_*ik,j*_ be the read count of taxon *j* (*j* = 1, …, *J*) in sample *i* (*i* = 1, …, *n*) obtained from data source *k* (*k* = 1, 2, 16S or SMS), and let *Z*_*i*_ be a vector of covariates excluding the intercept. To fit the LOCOM model simultaneously to 16S and SMS data, we choose one taxon (without loss of generality, taxon *J*) that has the largest mean relative abundance across both data sources to be the reference taxon (which does not need to be a null taxon), and compare each of the remaining taxa to the reference taxon using logistic regressions. Because the most abundant taxa can usually be effectively captured by both 16S and SMS, our selection criterion results in a reference taxon that is among the top abundant taxa in each data source. We define *p*_*ik,j*_ to be the expected value of the relative abundance in data source *k* and relate this relative abundance to the true relative abundance *π*_*i,j*_, which is unrelated to any data source, by log(*p*_*ik,j*_) = 𝕀 (*k* = 1)*γ*_1,*j*_ + 𝕀 (*k* = 2)*γ*_2,*j*_ +log(*π*_*i,j*_) + *α*_*ik*_, where *γ*_1,*j*_ and *γ*_2*j*_ are source-specific bias factors and *α*_*i,k*_ is the normalization factor that ensures 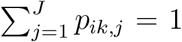. Following [30], we assume 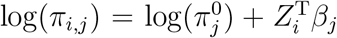, where *β*_*j*_ contains the effect sizes and 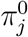 contains the baseline relative abundances. When comparing taxon *j* to the reference taxon *J*, we define *μ*_*ik,j*_ = *p*_*ik,j*_*/*(*p*_*ik,j*_ + *p*_*ik,J*_) to obtain a standard logistic regression

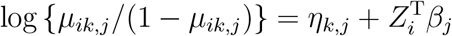

for each *j* (*j* = 1, 2, …, *J −* 1). The intercept *η*_*k,j*_ includes the taxon- and source-specific bias information, and for this reason is considered as a free parameter for each *j, k*.

To combine data from both sources for inference on the *β*_*j*_s, we solve the estimating equation (EE)

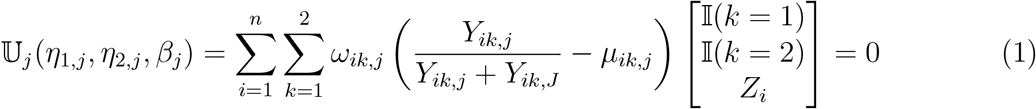

or the bias-corrected estimating equation (derived in Supplementary Materials) when taxon *j* has sparse count data, where *ω*_*ik,j*_ is a weight for the measurement of sample *i* from data source *k*. The EE of LOCOM is a special case of Equation (1) when *ω*_*ik,j*_ is set to *Y*_*ik,j*_ + *Y*_*ik,J*_ for one data source and *ω*_*ik,j*_ = 0 for the other.

When *ω*_*ik,j*_ = *Y*_*ik,j*_ + *Y*_*ik,J*_, the summation in Equation (1) is at the “count” level, and measurements with small *Y*_*ik,j*_ and *Y*_*ik,J*_ tend to contribute less. These weights are optimal when there is minimal overdispersion in the count data (as the equation becomes a score equation). Additionally, these weights are reasonable for a taxon when the count data from one data source are very sparse (i.e., mostly zeros) while the data from the other source are non-sparse. However, as taxa count data are frequently overdispersed, the amount of information quickly reaches a plateau as long as there is moderate coverage of reads. In this case, it is sensible to use *ω*_*ik,j*_ = 1, which puts the summation in Equation (1) at the “relative abundance” level and treats data from the two data sources equally. These weights are particularly important when the library sizes of SMS are orders of magnitude (e.g., 10 times) higher than that of 16S (Table S1), to prevent the SMS data from dominating the results. Since it is unknown a priori as for which weighting scheme works better for which taxa, Com-2seq fits (1) using both weights and then forms an omnibus test that combines both results. When a taxon has only been observed in one data source, Com-2seq only uses weights *ω*_*ik,j*_ = *Y*_*ik,j*_ + *Y*_*ik,J*_ and Equation (1) reduces to the LOCOM EE (Figure 1, right). When only relative abundance data are available, Com-2seq only uses weights *ω*_*ik,j*_ = 1.

Because there is no a priori knowledge about whether the reference taxon is null or causal (i.e., associated with the trait), Com-2seq follows LOCOM to reduce the dependence on the choice of reference taxon by testing each *β*_*j*_ against their median, which represents a null reference when assuming that more than half of the taxa are null taxa. Like LOCOM, Com-2seq bases inference on permutation using Potter’s method [31]. Care should be taken to preserve the sample structure when generating permutation replicates. The permutation procedure should be stratified to restrict the shuffling of trait residuals in one stratum of samples having data from both sources, one stratum of samples having data from 16S only, and one stratum of samples having data from SMS only (Figure 1, right). Finally, for testing the community-level (i.e., global) hypothesis that there are no differentially abundant taxa in the community, the Harmonic Mean (HM) of the taxon-specific *p*-values is used as the test statistic and assessed for significance using the permutation replicates generated above.

### Filtering out rare taxa in Com-2seq

We develop a filter to exclude very rare taxa that can jeopardize the validity of Com-2seq. Recall that the filter in LOCOM removes rare taxa present in fewer than 20% of samples. In Com-2seq, a sample with a non-zero count from either data source contributes to the 3rd equation of Equation (1), making the taxon analyzable if more than 20% of such samples exist. In addition, we keep the taxon as long as it passes the LOCOM filter applied to either 16S or SMS data, since the LOCOM filter works well for the “worse” case when the data from one source consist entirely of zeros.

### Alternative approaches Com-count and Com-p

Com-count first creates an “augmented” table consisting of the union of samples and the union of taxa across the two data sources, then pools (i.e., sums up) the read counts for cells that have both 16S and SMS data, copies the read counts for cells that have one data source, and fills in zeros for cells that have no data source. Then, LOCOM is applied to the augmented table. The zero-filling strategy may create spurious associations, when samples filled with zeros have a different trait distribution from the others. Moreover, the results of Com-count tend to be dominated by the data source that has much larger library sizes. Lastly, the LOCOM filter applied to the “augmented” table is more stringent than the Com-2seq filter above.

Com-p applies LOCOM separately to the two taxa count tables, and then combines the two *p*-values at each overlapping taxon into one *p*-value using the Cauchy *p*-value combination method [32]. The resulting *p*-values, along with *p*-values from LOCOM for non-overlapping taxa, are used to detect differentially abundant taxa after multiple test correction by the Benjamini-Hochberg method [33] and also combined into one *p*-value for testing the global hypothesis (also by the Cauchy method). At each taxon, the two *p*-values are expected to be correlated since they are based on at least some overlapping samples; at the global level, the *p*-values across interacting taxa may also be correlated. This motivated us to adopt the Cauchy [32] or HM [34] combination method, which accounts for such correlations; we chose Cauchy over HM because HM led to inflated error rate in our simulations (results not shown). Note that the HM method here produces *p*-values based on asymptotic theories, whereas Com-2seq assesses the significance of the HM statistics via permutation. Com-p is inherently unable to produce an overall *p*-value that is more significant than the most significant individual *p*-value, while Com-2seq can achieve this by combining the two data sources at the more appropriate taxon-count and relative-abundance levels. Additionally, Com-p combines *p*-values without considering directions of association in the two data sources, while Com-2seq is able to strengthen the signals if they are consistent across the two data sources and disregard the ones that are contradictory. Furthermore, taxa that fail the LOCOM filter applied to each taxa count table are completely missed by Com-p, but they still have a good chance of passing the Com-2seq filter (which is essentially based on the pooled data), especially when the sample overlap is substantial.

### Simulation settings

Our simulations are based on data on 856 taxa of the upper-respiratory-tract (URT) microbiome by Charlson et al. [35]. We fixed the sample size to 100 unless otherwise specified and considered a binary trait *T*_*i*_ throughout the simulations. In some cases, we also simulated a continuous confounder *C*_*i*_ by drawing values from *U* [*−*1, 1] for samples with *T*_*i*_ = 0 and from *U* [0, 2] for those with *T*_*i*_ = 1. We used the two sets of causal taxa employed in [23], M1 and M2, which are a random sample of 20 taxa with mean relative abundances greater than 0.005 and the five most abundant taxa, respectively. In addition, we considered a set of rare causal taxa by randomly sampling 50 taxa with mean relative abundances between 0.0005 and 0.001; we refer to this set as M3. When a confounder was present, we randomly sampled 5 taxa with mean relative abundances greater than 0.005 to be associated with the confounder.

We assumed that the 856 taxa form the complete set of underlying taxa in the community and generated bias factors *γ*_1,*j*_ and *γ*_2,*j*_ for taxon *j* from 16S and SMS, respectively. We set *γ*_1,*j*_ and *γ*_2,*j*_ to a very small value of *−*5 to create missingness for specific taxa in each data source. Specifically, in M1 (M2 and M3), we selected two sets of five (two and five) non-overlapping causal taxa to be missing in 16S and SMS, respectively. Additionally, we sampled 20% of non-causal taxa to be missing in each data source. We set *γ*_1,*j*_ and *γ*_2,*j*_ for the most abundant taxon to 1, reflecting the efficiency of both sequencing methods for capturing this taxon. For all other taxa, we independently drew *γ*_1,*j*_ and *γ*_2,*j*_ from *N* (0, 0.5^2^).

We then simulated read count data for the 856 taxa in the two data sources, taking into account the effects of the trait and confounder as well as the influences of bias factors. First, we drew the baseline relative abundances 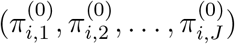 of all taxa for each sample from *Dirichlet*(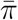, *θ*), where 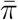 contains the mean relative abundances and *θ* is the overdispersion parameter estimated from fitting the Dirichlet-Multinomial (DM) model to the URT data. The overdispersion parameter *θ* controls the sample heterogeneity in baseline relative abundances, not including the variability in read count data. Thus, we set *θ* to 0.01, which is half of the overall overdispersion (0.02) estimated from the URT data. Then, we formed the expected value of the observed relative abundances obtained from the *k*th data source, *p*_*ik,j*_, by spiking in the causal taxa and confounder-associated taxa, then imposing bias factors on all taxa, and finally normalizing the relative abundances to have a sum of 1, resulting in the following equation:

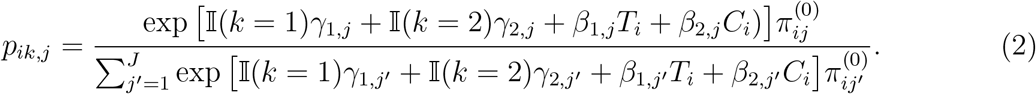

Here, *β*_1,*j*_ = 0 for null taxa, and *β*_2,*j*_ = 0 for confounder-independent taxa. For simplicity, we set *β*_1,*j*_ = *β* for all causal taxa, which is referred to as the *effect size*, and we fixed *β*_2,*j*_ = log(1.5) for all confounder-associated taxa. Subsequently, we generated the read count data for sample *i* in data source *k* from the DM distribution with mean (*p*_*ik*,1_, *p*_*ik*,2_, …, *p*_*ik,J*_), overdispersion parameter *τ*_*k*_, and library size drawn from *N* (*ν*_1_, (*ν*_1_*/*3)^2^) and *N* (*ν*_2_, (*ν*_2_*/*3)^2^) for 16S and SMS, respectively, with left truncation at 2,000. We fixed *ν*_1_ = 10, 000 and varied *ν*_2_ to achieve the depth ratio of *ν*_1_:*ν*_2_ = 1:1 or 1:10. The overdispersion parameter *τ*_*k*_ controls the deviation of the observed relative abundances from their expected values *p*_*ik,j*_ in the *k*th data source. We set *τ*_1_ = *τ*_2_ = *τ* without loss of generality and varied *τ* between 0.01 and 0.001, which correspond to the large and small deviation as seen in the Dietary and ORIGINS data, respectively.

In the complete-overlap case, we generated taxa count data from both sources for all 100 samples. In the partial-overlap case, we collected data from both sources for 40 samples (15 cases and 25 controls), from 16S only for 40 samples (30 cases and 10 controls), and from SMS only for 20 samples (5 cases and 15 controls), which resulted in a total of 100 samples, 50 cases and 50 controls, and varying case-control ratios across the three strata of samples.

We applied Com-2seq, Com-count, and Com-p to perform integrative analysis of 16S and SMS data, using the LOCOM results from analyzing individual datasets as benchmarks. We evaluated the sensitivity and empirical FDR of each method for testing individual taxa at the nominal FDR level of 20%, and the type I error and power of each method for testing the global association at the nominal level of 0.05. The type I error results were based on 10,000 replicates of simulated data, while all other results were derived from 1,000 replicates.

To confirm that the simulated data captured the important features of the ORIGINS data shown in Figure 2 and the Dietary data shown in Figure 3, we present scatter plots for the simulated data based on the sample size (*n* = 152) of the ORIGINS study and *τ* = 0.001 in Figure S6, and scatter plots for the simulated data based on the sample size (*n* = 76) of the Dietary study and *τ* = 0.01 in Figure S7. We observe that the range of observed relative abundances at each taxon as governed by the overdispersion parameter *θ*, the deviation of the fitted line from the 45^*°*^ reference line as determined by the bias factors *γ*_1,*j*_ and *γ*_2,*j*_, and the deviation of individual data points from the fitted line as controlled by the overdispersion parameter *τ*, all resemble those of the real data.

## Acknowledgments

This research was supported by the National Institutes of Health award R01GM141074 (Hu, Satten), and the Cancer Prevention and Research Institute of Texas (CPRIT) Rising Stars Award RR200056 (Fedirko).

## Notes

### Competing Interest Statement

The authors have declared no competing interest.

