## Supplementary Materials for "Integrative analysis of microbial 16S gene and shotgun metagenomic sequencing data improves statistical efficiency"

### Bias correction for generalized estimating equations

To simplify, we omit the taxon index $j$ and use the notation $\mathbb{U} = \mathbb{U}_j$, $\theta = \left(\eta_{1,j}, \eta_{2,j}, \beta_j^{\mathrm{T}}\right)^{\mathrm{T}}$, $X_{ik} = \left(\mathbb{I}(k=1), \mathbb{I}(k=2), Z_i^{\mathrm{T}}\right)^{\mathrm{T}}$, and $M_{ik} = Y_{ik,j} + Y_{ik,J}$. With this notation, we rewrite the estimation equation (1) as follows:

$$\mathbb{U}(\theta) = \sum_{i=1,\ldots n, k=1,2} \omega_{ik} \left( \frac{Y_{ik}}{M_{ik}} - \mu_{ik} \right) X_{ik} = 0. \tag{S1}$$

Let $\widehat{\theta}$ be the estimate of $\theta$ that solves Equation (S1). By taking a Taylor series expansion up to the 2nd order and using $\otimes$ to denote outer product, we obtain

$$\mathbb{U}(\widehat{\theta}) = 0 \approx \mathbb{U}(\theta) + \mathbb{J}(\theta)(\widehat{\theta} - \theta) + \frac{1}{2}(\widehat{\theta} - \theta)^{\mathrm{T}}\mathbb{K}(\theta)(\widehat{\theta} - \theta), \tag{S2}$$

where

$$\mathbb{J}(\theta) = \frac{\partial \mathbb{U}(\theta)}{\partial \theta} = -\sum_{i,k} \omega_{ik}\mu_{ik}\big(1 - \mu_{ik}\big) X_{ik} \otimes X_{ik}$$

and

$$\mathbb{K}(\theta) = \frac{\partial \mathbb{J}(\theta)}{\partial \theta} = -\sum_{i,k} \omega_{ik}\mu_{ik}\big(1 - \mu_{ik}\big)\big(1 - 2\mu_{ik}\big) X_{ik} \otimes X_{ik} \otimes X_{ik}.$$

We calculate the expected value of the right hand side of (S2), taking into account that $\mathrm{E}\big[\mathbb{U}(\theta)\big] = 0$ and that $\mathbb{J}(\theta)$ and $\mathbb{K}(\theta)$ are not functions of the data $Y_{ik}$, to obtain

$$0 = \mathbb{J}(\theta)\mathrm{E}(\widehat{\theta} - \theta) + \frac{1}{2}\mathrm{E}\Big\{(\widehat{\theta} - \theta)^{\mathrm{T}}\mathbb{K}(\theta)(\widehat{\theta} - \theta)\Big\}.$$

Let $b(\theta) = \mathrm{E}(\widehat{\theta} - \theta)$ be the asymptotic bias in $\widehat{\theta}$ and $\Sigma(\theta) = \mathrm{E}(\widehat{\theta} - \theta)(\widehat{\theta} - \theta)^{\mathrm{T}}$ be the variance-covariance matrix of $\widehat{\theta}$. Following from Firth [36], we write the bias-corrected estimating equation as

$$\mathbb{U}^*(\theta) = \mathbb{U}(\theta) + \mathbb{J}(\theta)b(\theta) = \mathbb{U}(\theta) - \frac{1}{2}\mathrm{trace}\Big[\mathbb{K}(\theta)\Sigma(\theta)\Big]. \tag{S3}$$

When the weight $\omega_{ik} = M_{ik}$, $\mathbb{U}(\theta)$ in (S1) is the score function for read count data that follow the Binomial distribution. In this case, the model-based variance-covariance estimator

$-\mathbb{J}(\theta)^{-1}$ is a reasonable estimator for $\Sigma(\theta)$, as adopted by LOCOM. When $\omega_{ik} = 1$, $\mathbb{U}(\theta)$ is no longer a score function and we estimate $\Sigma(\theta)$ using the robust sandwich estimator

$$\mathbb{J}(\theta)^{-1} \left[ \sum_{i,k} \omega_{ik}^2 \left( \frac{Y_{ik}}{M_{ik}} - \mu_{ik} \right)^2 X_{ik} \otimes X_{ik} \right] \mathbb{J}(\theta)^{-1}.$$

The model-based estimator might not be consistent in the presence of overdispersion in the read count data, and the sandwich estimator may not perform well with finite samples. However, it is important to note that we use these estimators solely for the purpose of bias correction in $\widehat{\theta}$. In the end, we rely on a permutation procedure to make valid inference.

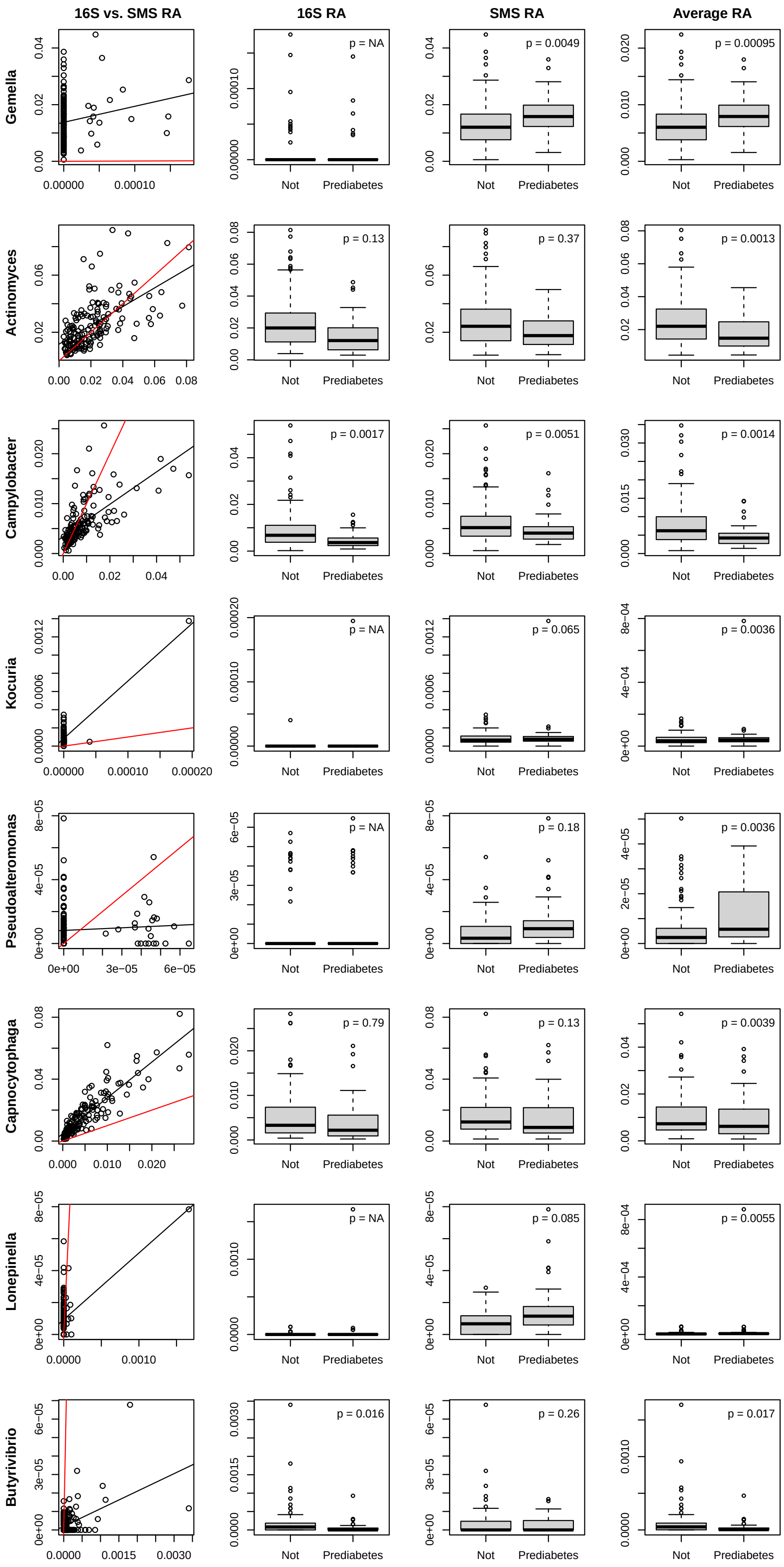


Figure S1: More information on the detected genera in the analysis of the ORIGINS data for overlapping samples and genera. In the first column, the scatter plots are the same as those in Figure 2. In the second and third columns, the $p$-values are from the analysis of 16S and SMS data, respectively, as also listed in Table S2. The last column shows the averages of observed relative abundances from 16S and SMS, along with the Com-2seq $p$-values.

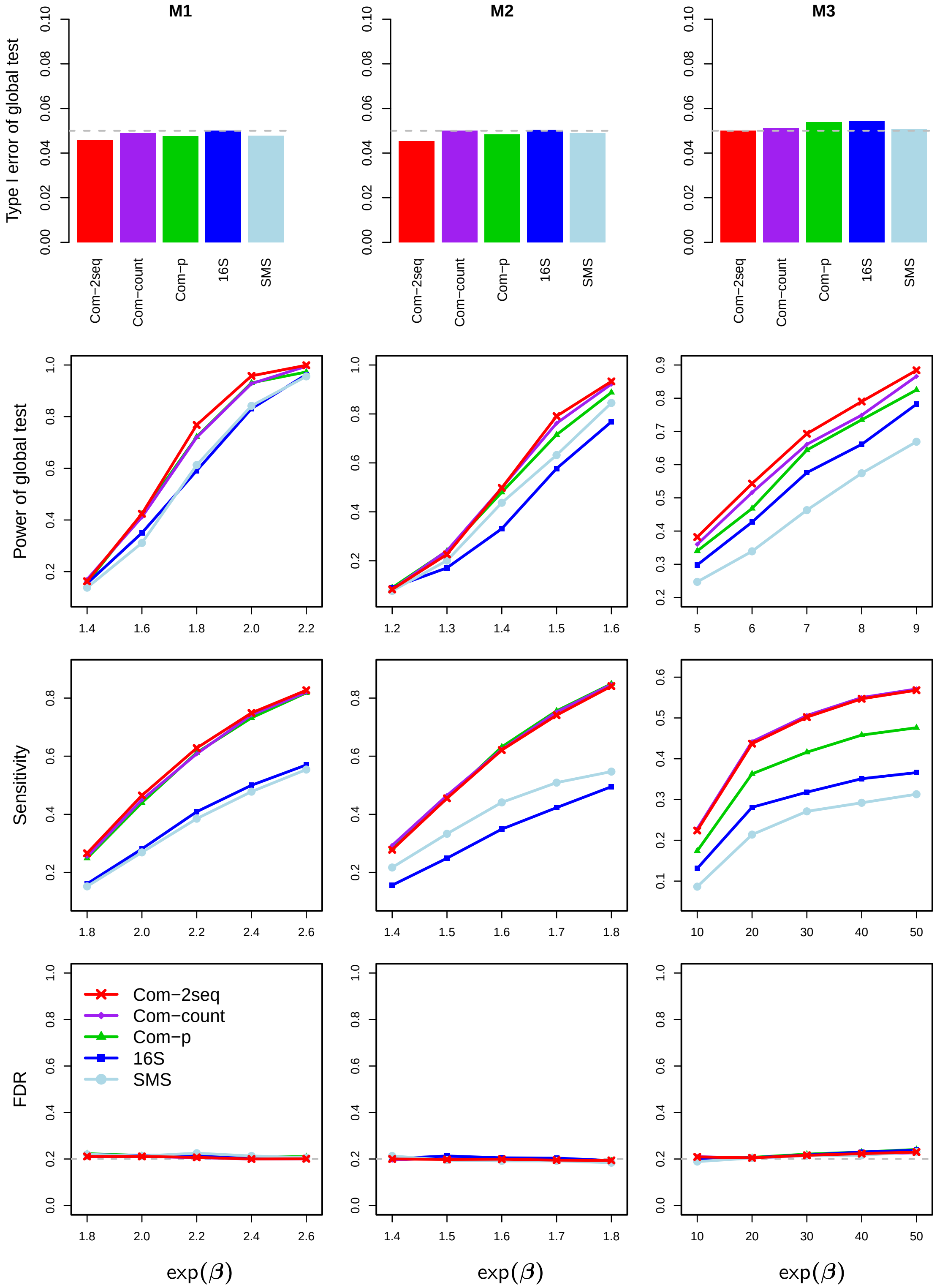


Figure S2: Results for data simulated with completely overlapping samples, overdispersion of $\tau = 0.01$, and depth ratio of 1:1.

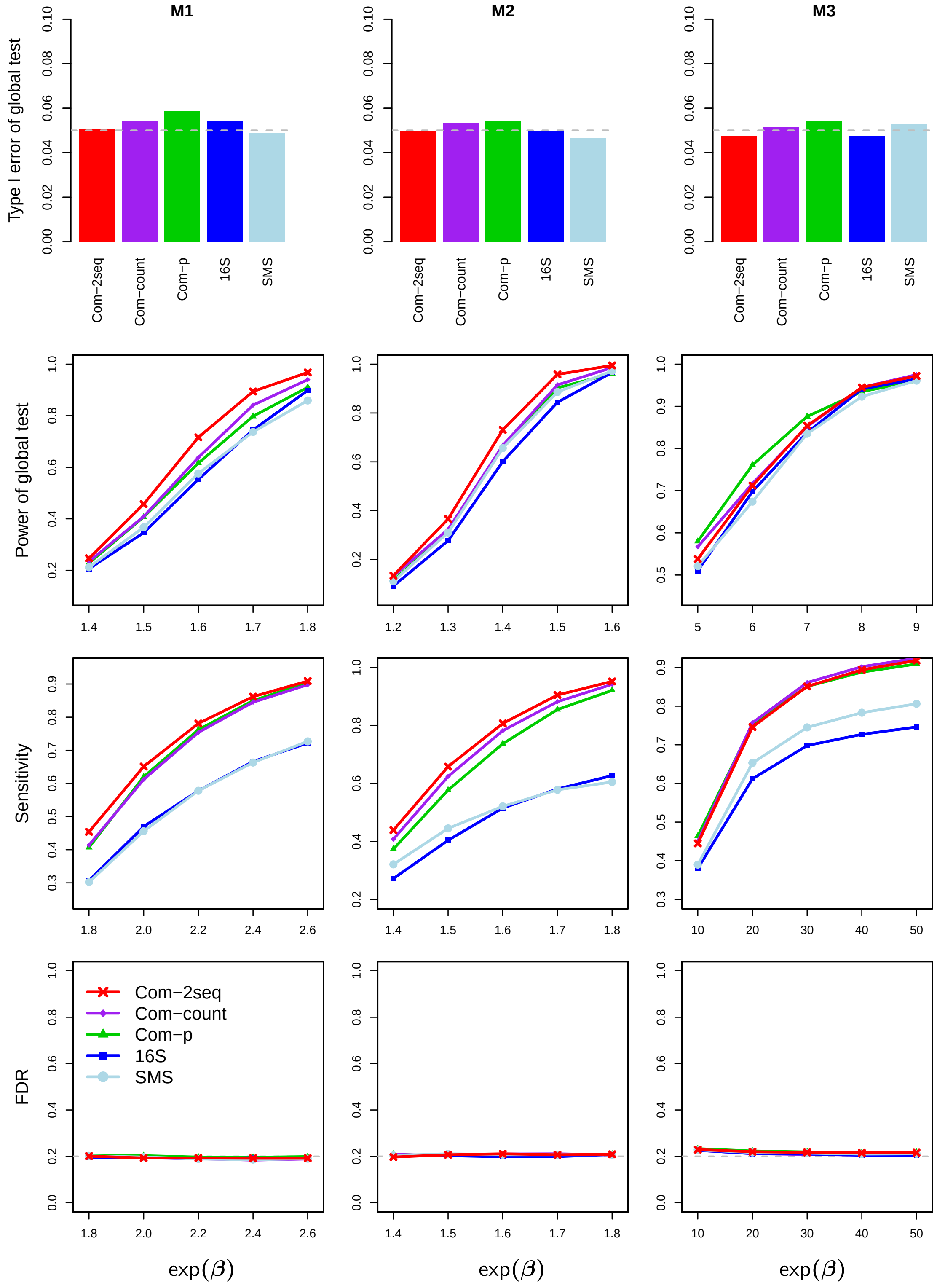


Figure S3: Results for data simulated with completely overlapping samples, overdispersion of $\tau = 0.001$, and depth ratio of 1:10.

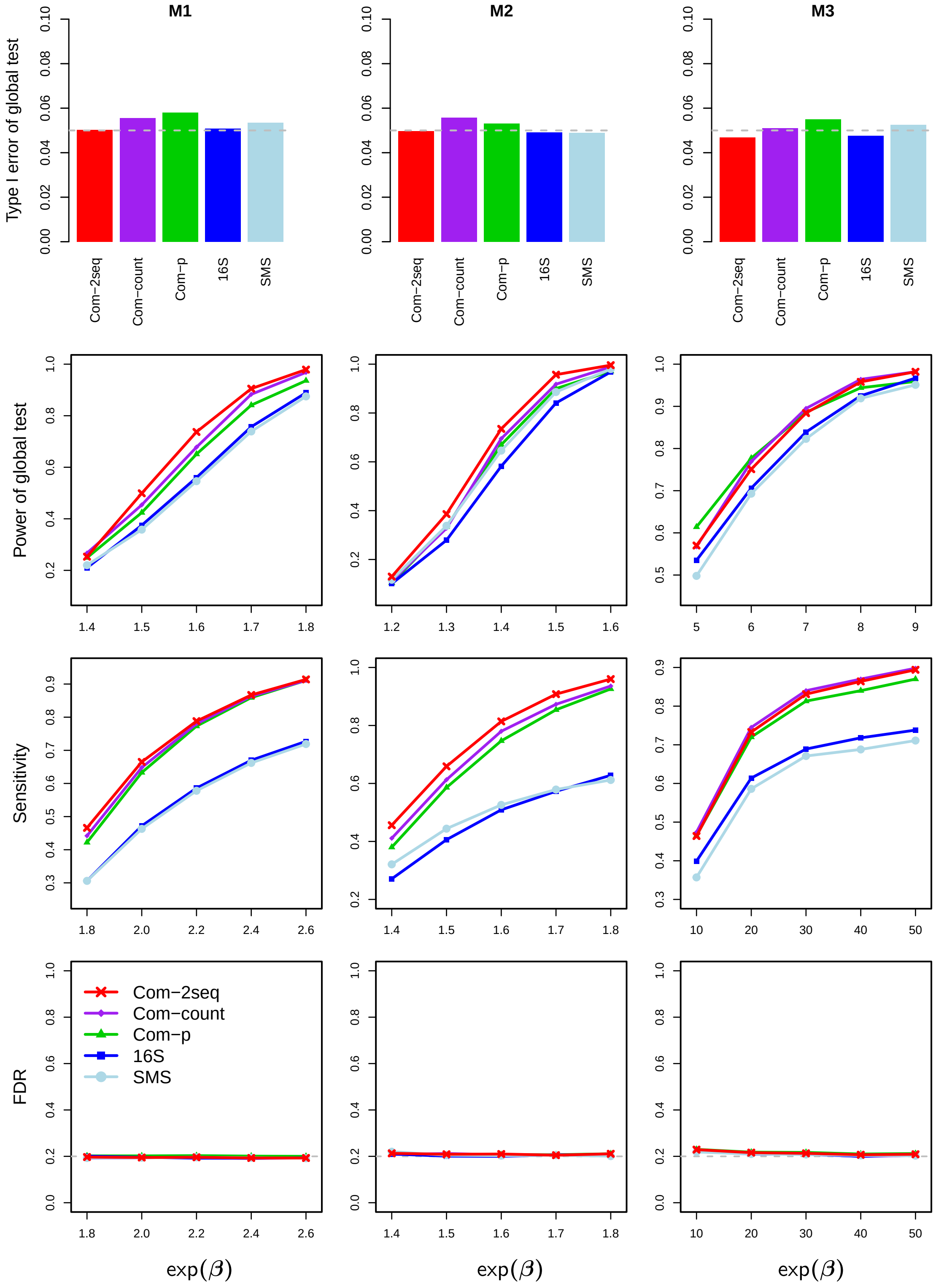


Figure S4: Results for data simulated with completely overlapping samples, overdispersion of $\tau = 0.001$, and depth ratio of 1:1.

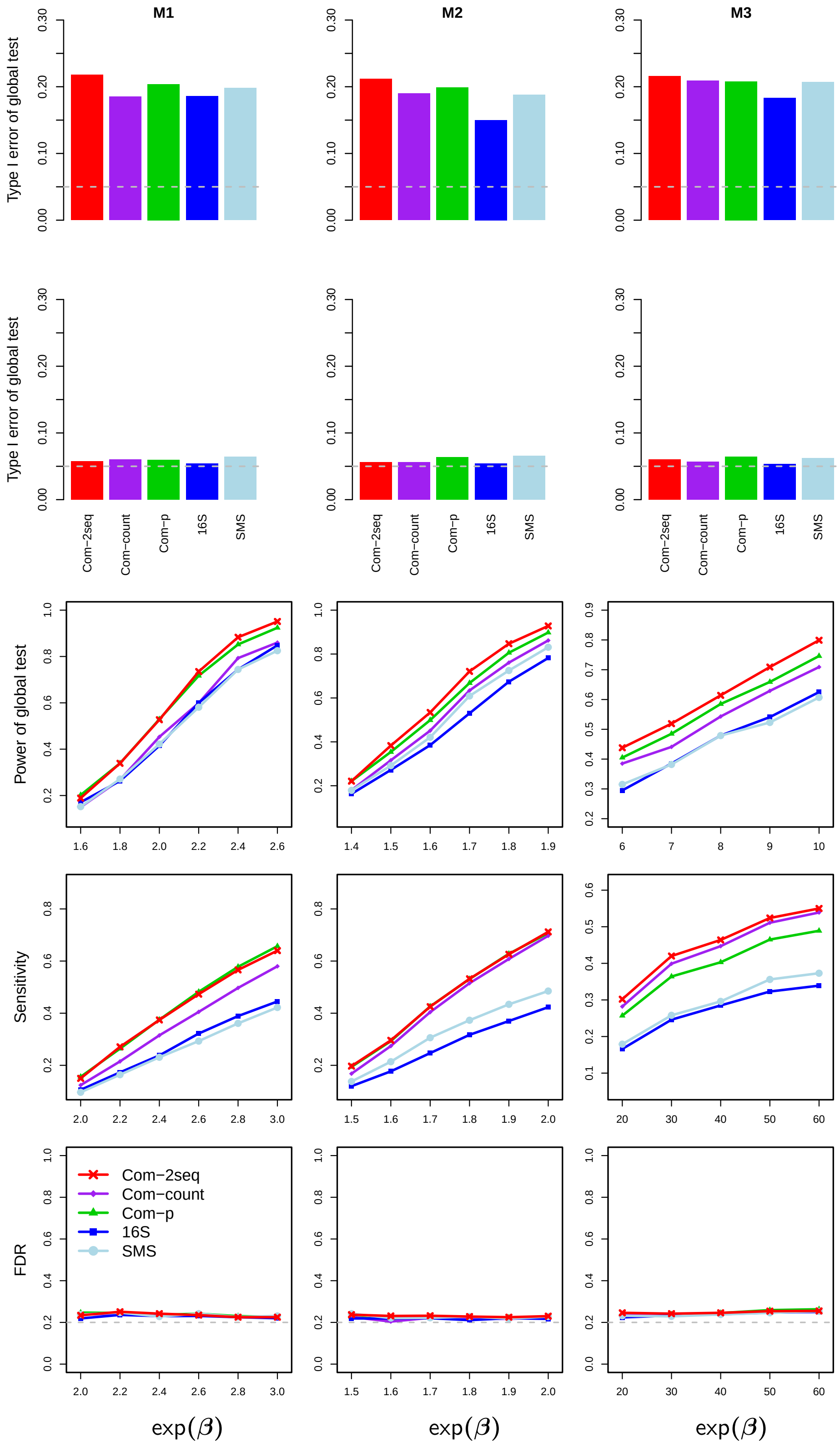


Figure S5: Results for data simulated with completely overlapping samples, a confounder, overdispersion of $\tau = 0.01$, and depth ratio of 1:10. The first row corresponds to global tests without adjusting for the confounder, while all other rows correspond to tests after adjustment for the confounder.

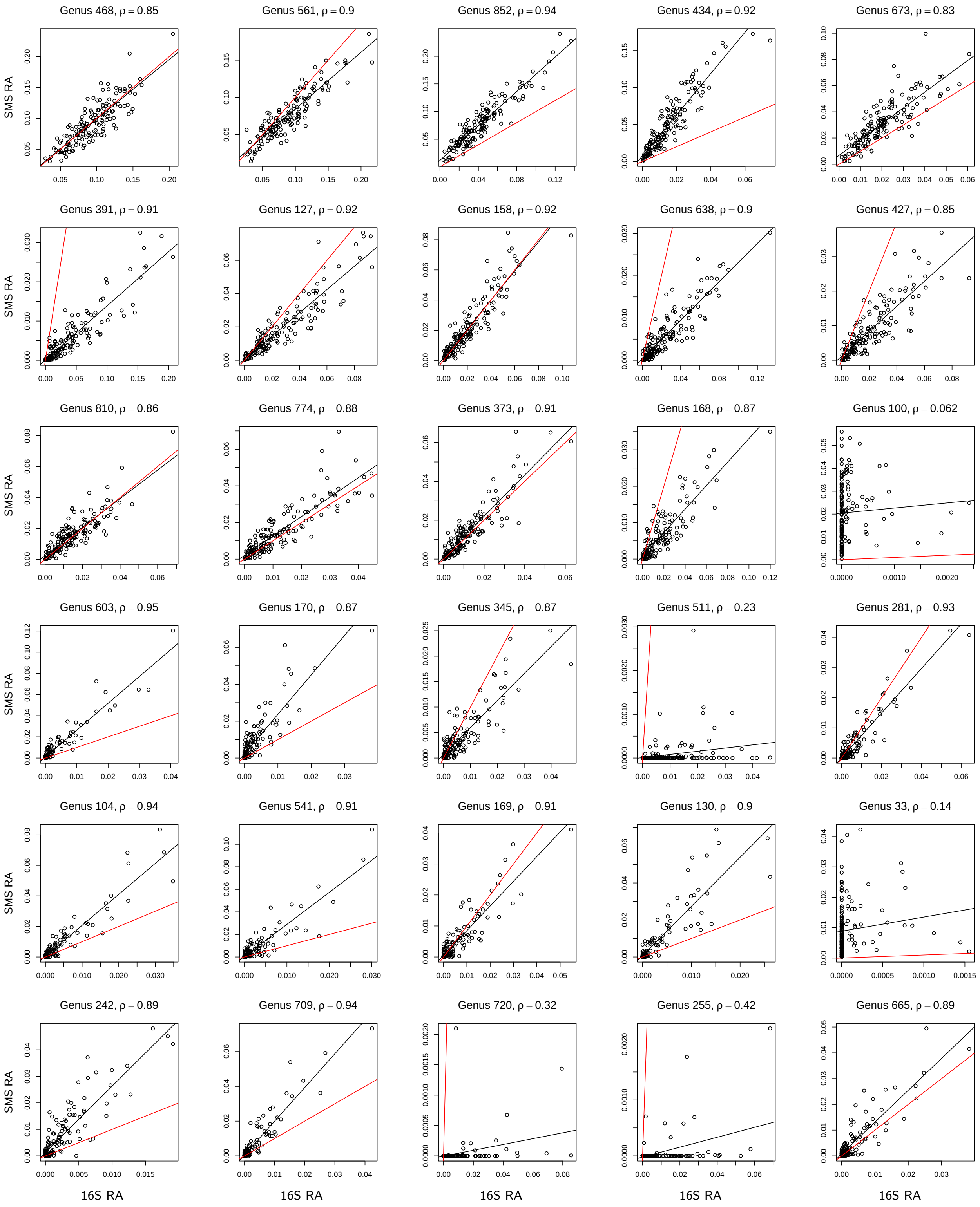


Figure S6: Scatter plot of observed relative abundances between 16S and SMS for the top 1–15 and 36–50 most abundant genera in the data simulated with 152 completely overlapping samples (the sample size as in Figure 2), overdispersion of $\tau = 0.001$, and depth ratio of 1:10. Find additional information in the caption of Figure 2.

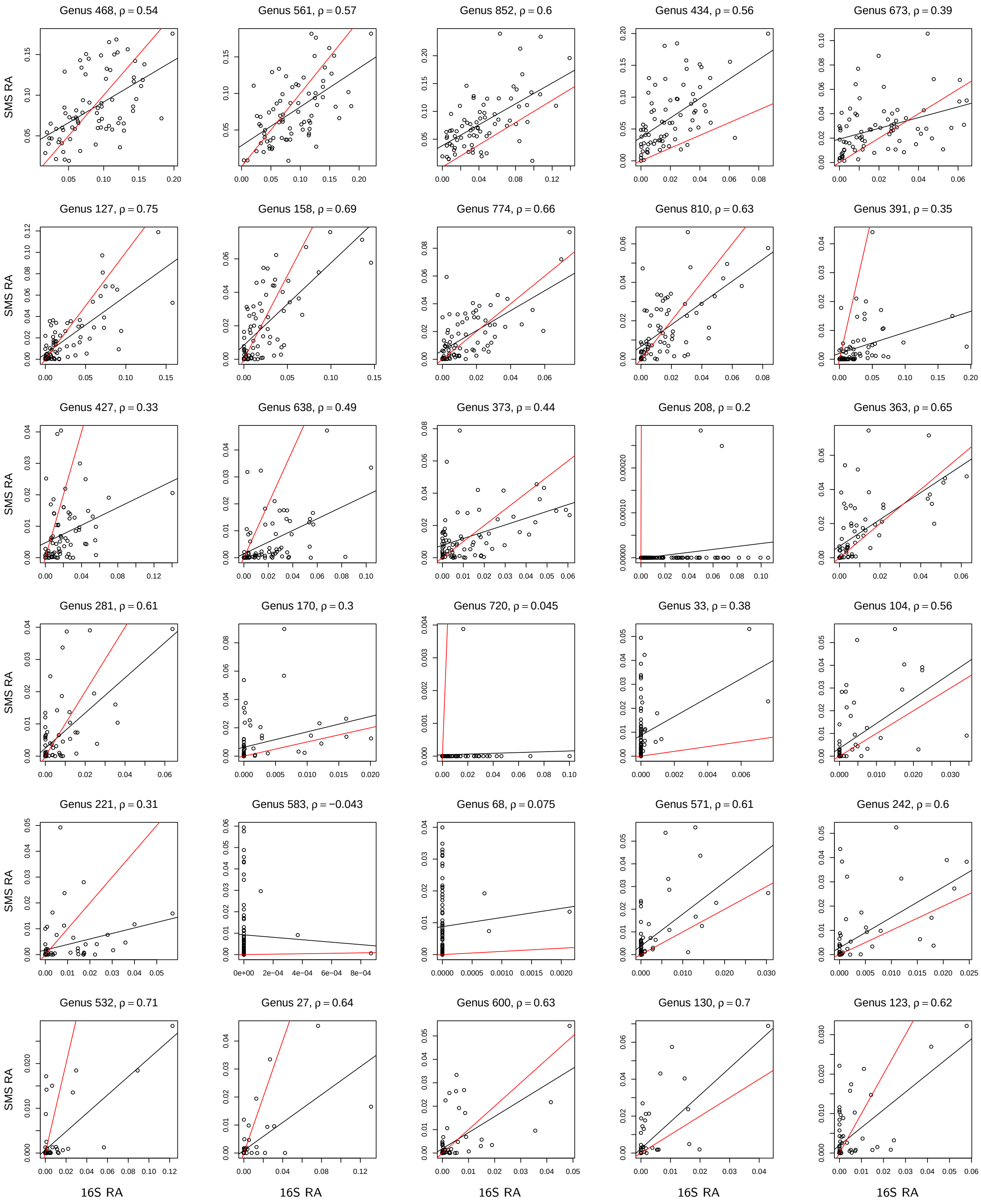


Figure S7: Scatter plot of observed relative abundances between 16S and SMS for the top 1–15 and 36–50 most abundant genera in the data simulated with 76 completely overlapping samples (the sample size as in Figure 3), overdispersion of $\tau = 0.01$, and depth ratio of 1:10. Find additional information in the caption of Figure 2.

Table S1: Studies in Qiita that have both 16S and SMS data.

| Qiita | 16S data | | | | SMS data | | | | Overlapping | | |
|---|---|---|---|---|---|---|---|---|---|---|---|
| Study ID | $n$ of sam | $n$ of OTU | $n$ of genus | Mean depth | $n$ of sam | $n$ of species | $n$ of genus | Mean depth | $n$ of sam | $n$ of genus | Depth ratio |
| 11479 | 591 | 3403 | 190 | 19487 | 907 | 2609 | 946 | 29085 | 518 | 106 | 1 : 1.5 |
| 10285 | 23 | 314 | 149 | 31004 | 12 | 839 | 372 | 51067 | 10 | 44 | 1 : 1.6 |
| *11808 | 271 | 884 | 234 | 26950 | 183 | 2147 | 756 | 176321 | 152 | 125 | 1 : 6.5 |
| 11841 | 257 | 2031 | 150 | 37370 | 54 | 1413 | 509 | 259332 | 53 | 55 | 1 : 6.9 |
| †11212 | 153 | 3933 | 524 | 17072 | 95 | 5084 | 1778 | 167100 | 76 | 236 | 1 : 9.8 |
| 11405 | 2096 | 3577 | 394 | 40001 | 1259 | 4791 | 1400 | 472294 | 1156 | 225 | 1 : 11.8 |
| 11896 | 87 | 696 | 135 | 45520 | 73 | 2542 | 834 | 565295 | 72 | 76 | 1 : 12.4 |
| 13114 | 237 | 9244 | 1357 | 15816 | 435 | 12535 | 2806 | 244093 | 144 | 564 | 1 : 15.4 |
| 11926 | 78 | 1007 | 115 | 18501 | 81 | 1945 | 755 | 301595 | 72 | 60 | 1 : 16.3 |
| 10394 | 1311 | 1195 | 300 | 44729 | 647 | 4042 | 1295 | 781702 | 599 | 142 | 1 : 17.5 |
| 11624 | 400 | 1744 | 365 | 25796 | 157 | 2382 | 750 | 456220 | 43 | 195 | 1 : 17.7 |
| 11484 | 58 | 4667 | 332 | 340647 | 150 | 5609 | 1745 | 11044763 | 17 | 174 | 1 : 32.4 |
| 11358 | 947 | 11743 | 1043 | 23753 | 40 | 2640 | 781 | 773496 | 40 | 290 | 1 : 32.6 |
| 11444 | 40 | 1408 | 123 | 40345 | 40 | 2392 | 656 | 1345059 | 40 | 74 | 1 : 33.3 |
| 11326 | 567 | 3785 | 240 | 17249 | 648 | 4394 | 1319 | 591237 | 118 | 138 | 1 : 34.3 |
| 13241 | 55 | 566 | 160 | 35410 | 96 | 3591 | 1034 | 2170485 | 55 | 86 | 1 : 61.3 |
| 11166 | 1312 | 20920 | 1480 | 92142 | 90 | 9011 | 2492 | 6222725 | 70 | 726 | 1 : 67.5 |
| 11149 | 24 | 500 | 111 | 29340 | 56 | 3114 | 933 | 3498073 | 20 | 72 | 1 : 119.2 |
| 13692 | 185 | 2750 | 262 | 45951 | 213 | 6451 | 1850 | 7349436 | 184 | 154 | 1 : 159.9 |
| 11673 | 138 | 2771 | 522 | 28994 | 67 | 3714 | 1100 | 4635809 | 64 | 231 | 1 : 159.9 |
| 2338 | 105 | 1256 | 428 | 93972 | 6 | 589 | 294 | 15856676 | 6 | 91 | 1 : 168.7 |
| 11549 | 40 | 714 | 997 | 12283 | 40 | 2882 | 823 | 4709741 | 38 | 61 | 1 : 383.4 |
| 10283 | 92 | 2462 | 212 | 54471 | 50 | 4781 | 1089 | 23960195 | 41 | 124 | 1 : 439.9 |
| 11546 | 306 | 2774 | 266 | 21684 | 367 | 7359 | 1495 | 33947551 | 285 | 147 | 1 : 1565.6 |

Note: "$n$"–number. "sam"–sample. *– ORIGINS study. †– Dietary study. The studies are ordered by depth ratio (i.e., ratio of 16S to SMS mean depths).

Table S2: $P$-value and adjusted $p$-value for the detected genera in the analysis of the ORIGINS data for overlapping samples and genera

| Method | *Gemella* | *Actino-myces* | *Campy-lobacter* | *Kocuria* | *Pseudo-alteromonas* | *Capnocy-tophaga* | *Lonepi-nella* | *Butyri-vibrio* |
|---|---|---|---|---|---|---|---|---|
| | | | | $p$-value | | | | |
| Com-2seq | 0.000947 | 0.00132 | 0.00137 | 0.00358 | 0.00363 | 0.00389 | 0.00553 | 0.0175 |
| Com-count | 0.000500 | 0.374 | 0.00090 | 0.0133 | 0.0501 | 0.248 | 0.0165 | 0.0005 |
| Com-p | 0.00491 | 0.201 | 0.00257 | 0.0647 | 0.183 | 0.347 | 0.0852 | 0.0300 |
| 16S | NA | 0.131 | 0.00172 | NA | NA | 0.794 | NA | 0.0157 |
| SMS | 0.00491 | 0.368 | 0.00509 | 0.0647 | 0.183 | 0.127 | 0.0852 | 0.261 |
| | | | | Adjusted $p$-value | | | | |
| Com-2seq | 0.0447 | 0.0447 | 0.0447 | 0.0636 | 0.0636 | 0.0636 | 0.0774 | 0.1900 |
| Com-count | 0.0245 | 0.7200 | 0.0294 | 0.2020 | 0.3510 | 0.6570 | 0.2020 | 0.0245 |
| Com-p | 0.236 | 0.682 | 0.236 | 0.415 | 0.652 | 0.818 | 0.481 | 0.288 |
| 16S | NA | 0.634 | 0.093 | NA | NA | 0.967 | NA | 0.212 |
| SMS | 0.229 | 0.862 | 0.229 | 0.531 | 0.660 | 0.660 | 0.565 | 0.730 |

Note: “NA” means that the genus failed to pass the LOCOM filter.

Table S3: $P$-value and adjusted $p$-value for the detected genera in the analysis of the full ORIGINS data

| Method | *Butyrivibrio* | *Gemella* | *Ignavigranum* | *Chelonobacter* |
|---|---|---|---|---|
| | | $p$-value | | |
| Com-2seq | 0.00042 | 0.00052 | 0.00014 | 0.189 |
| Com-p | 0.120 | 0.00058 | 0.00062 | 0.0015 |
| 16S | 0.0711 | NA | NA | 0.00075 |
| SMS | 0.313 | 0.00058 | 0.00062 | 0.727 |
| | | Adjusted $p$-value | | |
| Com-2seq | 0.0763 | 0.0763 | 0.0616 | 0.617 |
| Com-p | 0.499 | 0.136 | 0.136 | 0.191 |
| 16S | 0.465 | NA | NA | 0.0637 |
| SMS | 0.741 | 0.125 | 0.125 | 0.945 |

Note: See Note in Table S2.
